# To the canopy and beyond: Air samples reveal wind dispersal as a driver of ubiquitous protistan pathogen assembly in tree canopies

**DOI:** 10.1101/2020.11.30.405688

**Authors:** Robin-Tobias Jauss, Anne Nowack, Susanne Walden, Ronny Wolf, Stefan Schaffer, Barbara Schellbach, Michael Bonkowski, Martin Schlegel

**Affiliations:** University of Leipzig, Institute of Biology, Molecular Evolution & Animal Systematics, Talstraße 33, 04103 Leipzig; University of Cologne, Institute of Zoology, Terrestrial Ecology, Zülpicher Straße 47b, 50674 Köln; Max Planck Institute for Evolutionary Anthropology, Department of Evolutionary Genetics, Deutscher Platz 6, 04103 Leipzig; German Centre for Integrative Biodiversity Research (iDiv) Halle Jena Leipzig, Deutscher Platz 5e, 04103 Leipzig

**Keywords:** Airborne Microorganisms, Protists, Forest Ecosystems, Canopies, Pathogens

## Abstract

We analyzed air dispersal of the protistan phyla Cercozoa and Oomycota with an air sampler near the ground (~2 m) and in tree crowns (~25 m) of three tree species (oak, linden and ash) in a temperate floodplain forest in March (before leafing) and May (after leaf unfolding) with a cultivation-independent high throughput metabarcoding approach. Both, Cercozoa and Oomycota, contain important pathogens of forest trees and other vegetation. We found a high diversity of Cercozoa and Oomycota in air samples with 122 and 81 OTUs, respectively. Especially oomycetes showed a high temporal variation in beta diversity between both sampling dates. Differences in community composition between air samples in tree canopies and close to the ground were however negligible, and also tree species identity did not affect communities in air samples, indicating that the distribution of protistan propagules through the air was not spatially restricted in the forest ecosystem. OTUs of plant pathogens, whose host species that did not occur in the forest, demonstrate wind dispersal of propagules from outside the forest biome. Overall, our results lead to a better understanding of the stochastic processes of wind dispersal of protists and protistan pathogens, a prerequisite to understand the mechanisms of their community assembly in forest ecosystems.

**Importance:** Wind dispersal has been shown to play a crucial role in protistan community assembly. The protistan taxa Cercozoa and Oomycota contain important plant parasites with a major ecologic and economic impact. However, comprehensive assessments of cercozoan and oomycete diversity in forest air samples were lacking. Using a cultivation-independent high throughput metabarcoding approach, we analyzed cercozoan and oomycete air dispersal in forest floors and the canopy region – a potential filter for microbial propagules. Our study provides insights into the diversity and community assembly of protists within the air, contributing to a better understanding which factors drive the distribution of plant pathogens within forest ecosystems.

## 1. Introduction

The air is an effective means of long-distance propagation for a wide range of microbial organisms (Foissner & Hawksworth, 2009; Pepper & Dowd, 2009). The phyllosphere – and especially the crowns of trees – are the largest biological interface between the soil and the atmosphere (Ozanne et al., 2003; Ellwood & Foster, 2004), which therefore may act as a huge natural filter for airborne microbial propagules, including unicellular Eukaryotes (Protists). Within the paraphyletic taxon of protists, the group of Cercozoa (Rhizaria) are highly diverse in morphology and physiology and show a high functional and ecological variety (Bass et al., 2009; Harder et al., 2016). They dominate terrestrial habitats (Urich et al., 2008; Voss et al., 2019) and harbor important plant pathogens, such as the Endomyxa, which have recently been elevated from the Cercozoa into a separate phylum (Cavalier-Smith et al., 2018). Another protistan taxon, the Oomycetes (Stramenopiles), contain important parasites of forest trees, and many lineages produce caducous sporangia for dissemination (Goheen & Frankel, 2009; Robideau et al., 2011; Lang-Yona et al., 2018). With almost 800 described species, Oomycota are reported to have a broad distribution and a wide variety of ecological roles (Robideau et al., 2011; Thines, 2014; Judelson, 2017). Further, it is one of the eukaryotic groups with a great impact on ecosystems, as well as on economics and human health: the most famous species is *Phytophthora infestans*, which causes the potato blight. In the 1840s it led to the great famine in Ireland followed by massive emigration (Lara & Belbahri, 2011; Robideau et al., 2011).

Protists can be passively disseminated over long distances by viable propagules, mostly as resting stages (cysts), while some groups, especially pathogens with more complex life cycles, also form sporangia for dispersal (Cowling, 1994; Kageyama & Asano, 2009). Cysts are formed under unfavorable conditions, e.g. due to dryness, lack of food, or microbial antibiotics (Petz & Foissner, 1988; Adl & Gupta, 2006; Jousset et al., 2006), and it has been assumed that the cyst bank plays an important role for the resilience of protists and their functions in terrestrial environments (Geisen et al., 2017). Viable protist cysts can be retrieved from soils even after decades (Moon-van der Staay et al., 2006; Kageyama & Asano, 2009), leading to the long-standing question of how cosmopolitan protists are (Finlay, 2002; Fenchel & Finlay, 2004; Foissner, 2009).

Finlay et al. (2001) proposed that the spatial distribution of protistan propagules is influenced by several randomizing factors, such as soil particles dispersed by wind, convective transport, percolating rainwater, fog or animals. Rogerson and Detwiler (1999) determined that on average 0.25 cysts m^−3^are contained in the air depending on wind speed and time since last precipitation. Using a molecular approach, Genitsaris et al. (2014) came to generally similar conclusions, while they further detected operational taxonomic units (OTUs) with constant presence as well as OTUs exhibiting seasonal variation. High humidity increases the chance of survival of transported microbes and promotes their deposition (Fuzzi et al., 1997; Evans et al., 2019) and airborne microorganisms can be transported in fog droplets by atmospheric turbulence over long distances (Fuzzi et al., 1997; Amato et al., 2005).

Recently, Jauss et al. (2020) confirmed a ubiquitous distribution of Cercozoa and Oomycota in a floodplain forest, despite strong differences in community composition of different microhabitats related to differences in the relative abundance of taxa. This led to the conclusion that within forest ecosystems both cercozoans and oomycetes can colonize most habitats, in which they then however do not perform similarly well due to habitat filtering. One reason for this ubiquitous presence of these protists could be wind dispersal.

Here, we studied the air dispersal of Cercozoa, Endomyxa and Oomycota by a cultivation-independent high throughput metabarcoding approach to analyze protistan diversity in the air surrounding tree canopies and near the ground of a temperate floodplain forest, to gain a deeper insight into the mechanisms how protists and their pathogenic lineages are distributed in the environment. These examinations tackled three hypotheses: (1) Wind dispersal explains the ubiquitous presence of these protists in the floodplain forest. (2) There are differences in the distribution in the vertical plane as a strong discrepancy between canopy and ground habitats was previously described. (3) Temporal variation of wind dispersed propagules further drives the community and pathogen assembly in forest ecosystems.

## 2. Material and Methods

### 2.1. Sampling and DNA extraction

Air samples were taken in a temperate deciduous floodplain forest in the northwest of the city of Leipzig, Germany (51.3657 N, 12.3094 E) with a MicroBio MB2 Bioaerosol Sampler (Cantium Scientific, Dartford, UK) containing 1% agar plates. The samples were collected under defined conditions drawing 100 l/min of air for ten minutes in two strata: (1) near the ground (~2m) and (2) in ~25m height in tree canopies with the help of the Leipzig Canopy Crane (LCC) facility. Two samplings were carried out – one in March and one in May 2019. For each sampling, three tree species with three replicates each were chosen (*Quercus robur, Tilia cordata* and *Fraxinus excelsior*). As non-arboral control, samples were also taken on the crane tracks near the ground and at canopy height. Two plates per stratum and of each replicated tree species were collected, yielding 40 plates per sampling. After air sampling, the agar plates were taken out of the instrument, sealed with parafilm to prevent contaminations and frozen until the DNA was extracted with the DNeasy PowerSoil^®^ Kit according to the instructions supplied by the manufacturer. Weather conditions were tracked with a WebVIS data logger attached to the crane (Umweltanalytische Produkte GmbH, Ibbenbüren, Germany) (Table 1).

**Table 1:**
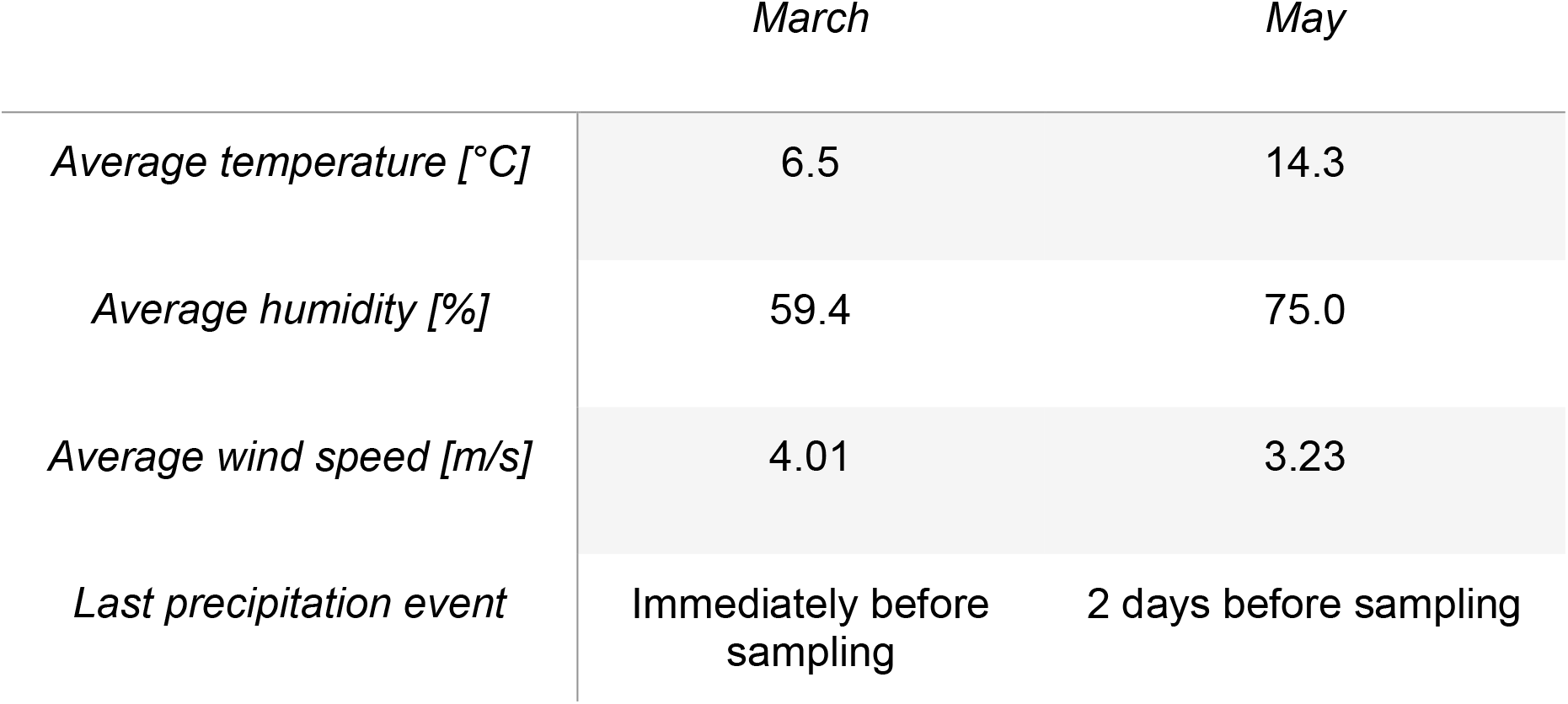
Weather conditions at the sampling days in March and May 2019.

|  | March | May |
| --- | --- | --- |
| Average temperature [°C] | 6.5 | 14.3 |
| Average humidity [%] | 59.4 | 75.0 |
| Average wind speed [m/s] | 4.01 | 3.23 |
| Last precipitation event | Immediately before sampling | 2 days before sampling |

### 2.2. PCR amplification and sequencing

DNA was amplified in duplicates with tagged oomycete- and cercozoan-specific primers (Fiore-Donno & Bonkowski, 2020; Jauss et al., 2020; Fiore-Donno et al., 2020) (Supplementary Tables 1-2). PCR-products were purified following the directions of the NucleoSpin^®^ PCR clean-up protocol. Afterwards, DNA concentrations were measured with the Qubit™4 fluorometer in combination with the Qubit™ dsDNA HS Assay Kit. For consecutive Illumina MiSeq Sequencing, a library was prepared following the Meyer and Kircher (2010) protocol. DNA concentrations were checked repeatedly before and after Illumina sequencing by utilization of DNA chips analyzed with the Agilent 2100 Bioanalyzer. Between the steps of library preparation, reaction clean-up was performed with the AMPure XP System using carboxyl coated magnetic beads (SPRI beads). Subsequent steps and the Illumina MiSeq sequencing itself were performed by the sequencing team of the Max Planck Institute for Evolutionary Anthropology in Leipzig, Germany.

### 2.3. Sequence processing and statistical analyses

Bioinformatic and statistical analyses followed the pipeline described in Jauss et al. (2020). Briefly, resulting reads were merged and clustered into operational taxonomic units (OTUs) using a custom pipeline utilizing cutadapt v1.18 (Martin, 2011), Swarm v2.2.2 (Mahé et al., 2015) and VSEARCH v2.10.3 (Rognes et al., 2016). OTUs were then annotated using NCBI’s non-redundant nucleotide database and the Protist Ribosomal Reference Database (Guillou et al., 2013) for oomycete and cercozoan OTUs, respectively (Supplementary Tables 3-4). OTUs resembling non-oomycete or non-cercozoan sequences were excluded. Samples with less than 5 OTUs or with a sequencing depth lower than 20617 reads (Oomycota) and 16922 reads (Cercozoa) were omitted. Statistical analyses of alpha and beta diversity and final visualizations were performed in R v3.5.3 (R Core Team, 2019) with the packages vegan (Oksanen et al., 2019), ggplot2 (Wickham, 2016) and ggraph (Pedersen, 2020).

## 3. Results

### 3.1. Amplification, sequencing and bioinformatic pipeline

After DNA isolation, all oomycete samples were amplified successfully whereas nine out of 20 cercozoan samples had to be excluded due to the failure of successful amplification of duplicates. Further, samples containing less than 1ng/μl DNA were excluded from subsequent processing, as well as samples with a low sequencing depth (see 2.3), yielding 9 cercozoan samples from March and 4 from May, as well as 13 oomycete samples from March and 20 from May (Supplementary Table 5). Of cercozoan sequences, 94.4% could be merged with a mean length of 370±35 bp resulting in 122 OTUs in total. Of oomycete sequences, 92.6% of derived 300 bp long paired-end sequences could be merged and the mean fragment length accounted for 285±38 bp, which were finally clustered into 81 OTUs.

### 3.2. Alpha diversity

Neither species richness nor Shannon-diversity nor evenness of Cercozoa or Oomycota differed between tree species, ground and canopy or non-arboreal controls, although variation of canopy samples was much lower than of ground samples in Cercozoa (e.g. species richness CV_Ground_ = 42.3%, CV_Canopy_ = 5.6%). However, Shannon-diversity and evenness of both protistan groups and species richness of oomycetes were higher in May than in March, indicating that the tree foliage in May did not restrict protistan distribution (Figure 1).

**Figure 1:**
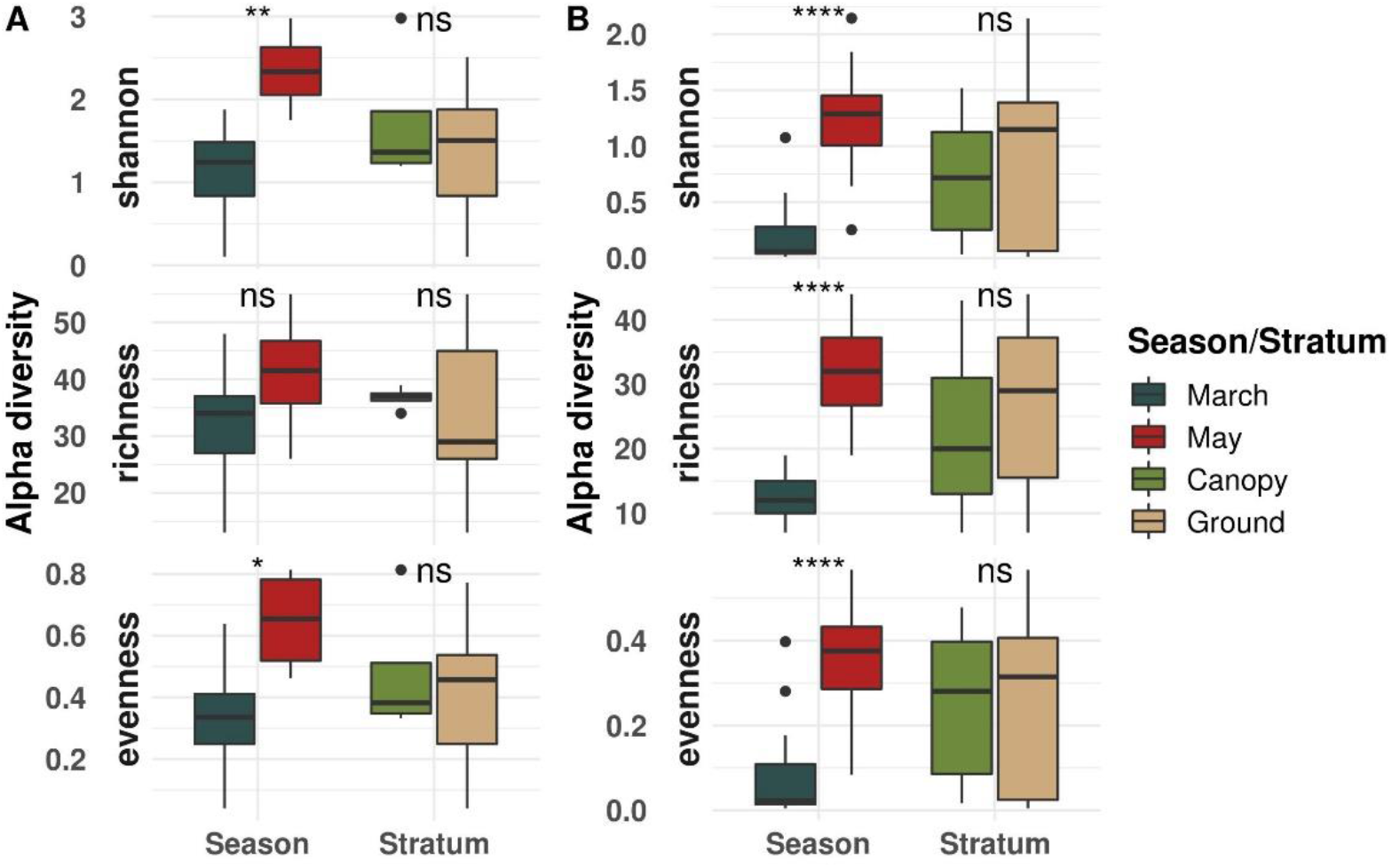
Boxplot of alpha diversity indices of cercozoan (A) and oomycete (B) samples. Pairwise comparisons of March and May samples and canopy and ground samples, respectively are shown. Significance was tested with Wilcoxon Sign test and is indicated by asterisks (ns = p>0.05, * = p<0.05, ** = p<0.01, *** = p<0.001, **** = p<0.0001).

### 3.3. Beta diversity

For Cercozoa, β-diversity of air samples did not differ between tree species (permANOVA R^2^=0.165, p=0.514), ground vs. canopy stratum (permANOVA R^2^=0.093, p=0.296) nor sampling season (permANOVA R^2^=0.101, p=0.168). However, variation of β-diversity was much lower in May compared to March, and lower in canopy samples compared to ground samples (Figure 2A). Oomycete communities differed between sampling seasons (permANOVA R^2^=0.170, p=0.001), but not between tree species (permANOVA R^2^=0.080, p=0.719) or the strata ground and canopy (permANOVA R^2^=0.037, p=0.259) (Figure 2B).

**Figure 2:**
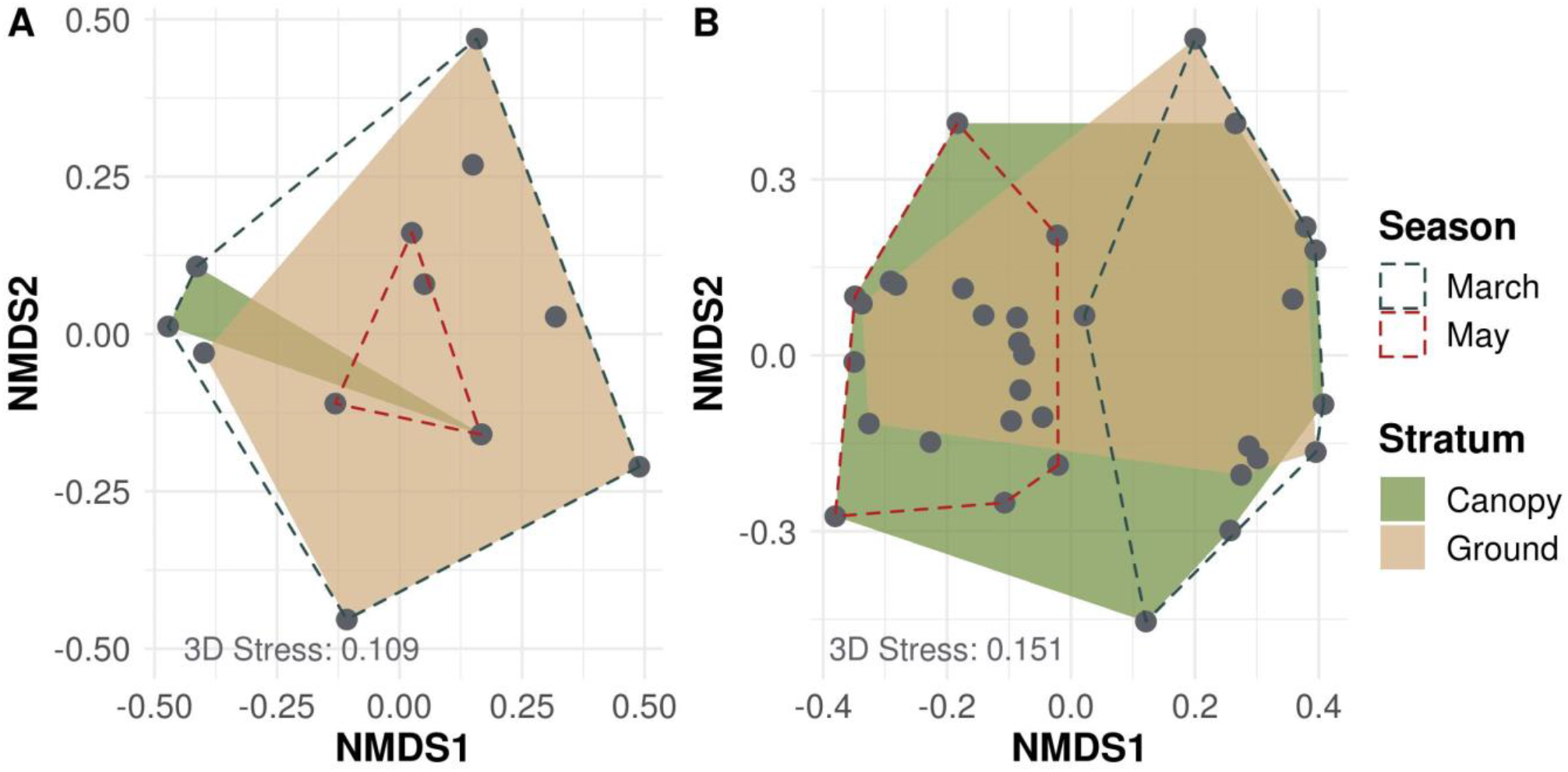
Non-metric multidimensional scaling (NMDS) plot of cercozoan (A) and oomycete (B) samples. Canopy and ground samples show a large overlap, while in oomycetes the March and May samples show a strong separation.

### 3.4. Taxonomic diversity

Cercozoan OTUs were dominated by the orders Cryomonadida and Glissomonadida, whereas the least abundant ones were Marimonadida and an unspecified order named Cercozoa_XX, comprising undescribed cercozoan lineages (Figure 3A). We detected no OTUs assigned to the plant parasitic group of Endomyxa. Oomycete OTUs were almost exclusively dominated by Peronosporales (Figure 3B), with only few members of the Pythiales, and the Albuginales being the least abundant order.

**Figure 3:**
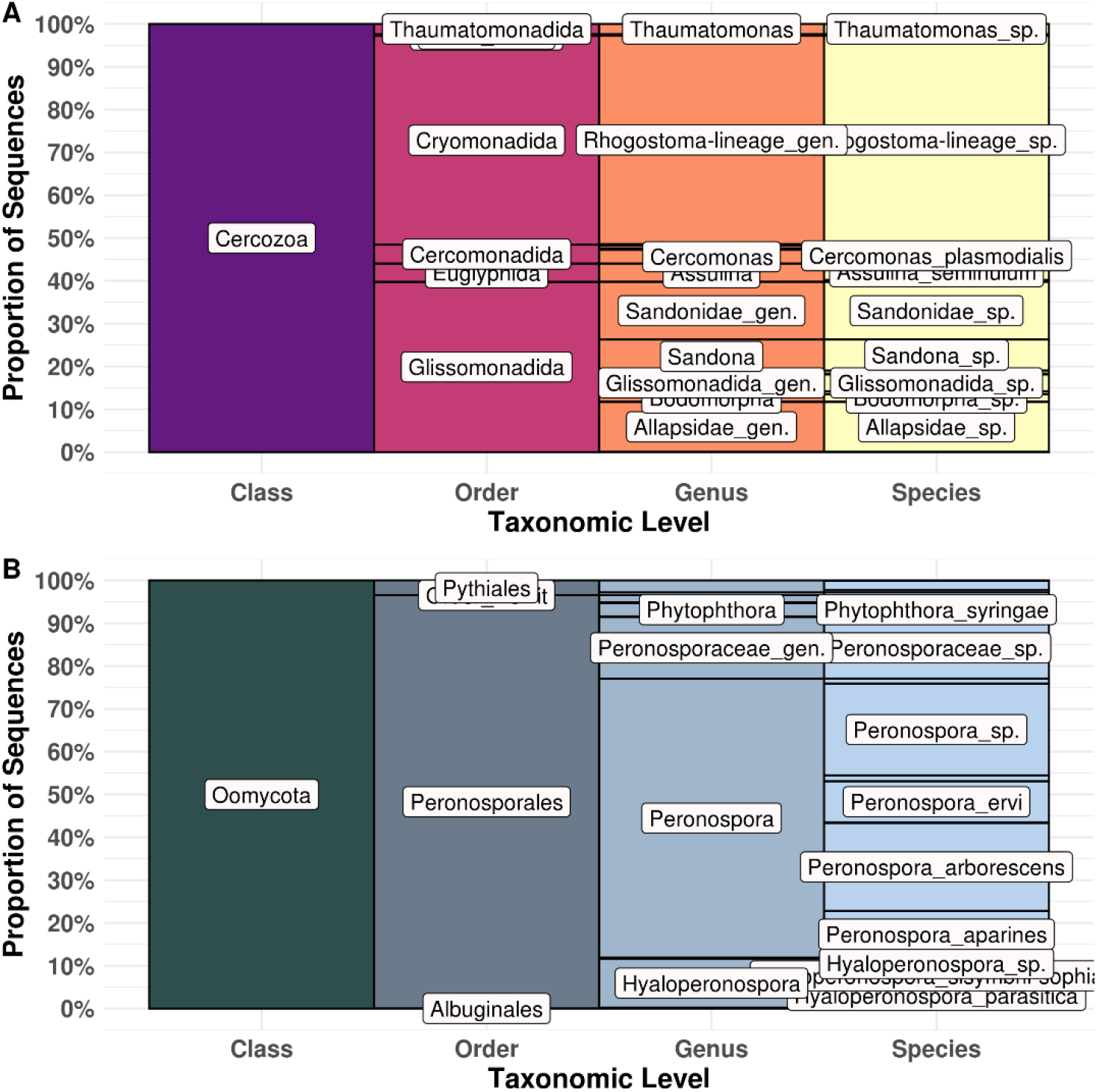
Taxonomic annotation of cercozoan (A) and oomycete (B) OTUs. Labels give the detected orders and the ten most abundant species with their corresponding genus.

The number of shared OTUs indicated a temporal variation in air dispersal of both protistan taxa (Figure 4), and dispersal of cercozoan OTUs varied also spatially between canopy and ground at the incidence level.

**Figure 4:**
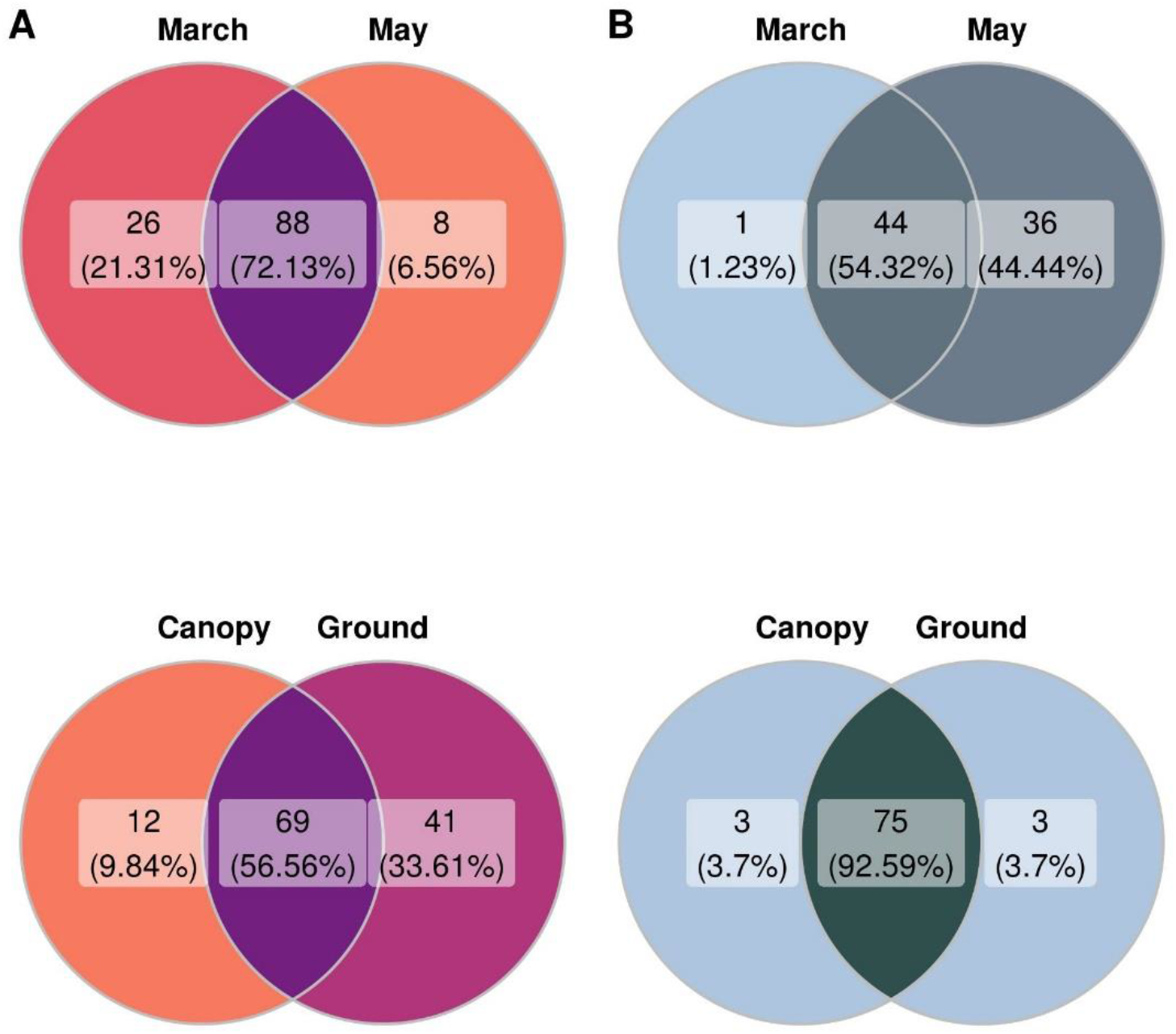
Venn diagram giving the number and proportion of shared OTUs between March and May samples and Canopy and Ground samples, respectively, for Cercozoa (A) and Oomycota (B).

Partitioning of the taxonomy into the two sampling seasons revealed similar patterns (Figure 5), yet, the cercozoan order Euglyphida was exclusively present in March samples, and oomycete Pythiales showed a higher abundance in March than in May.

**Figure 5:**
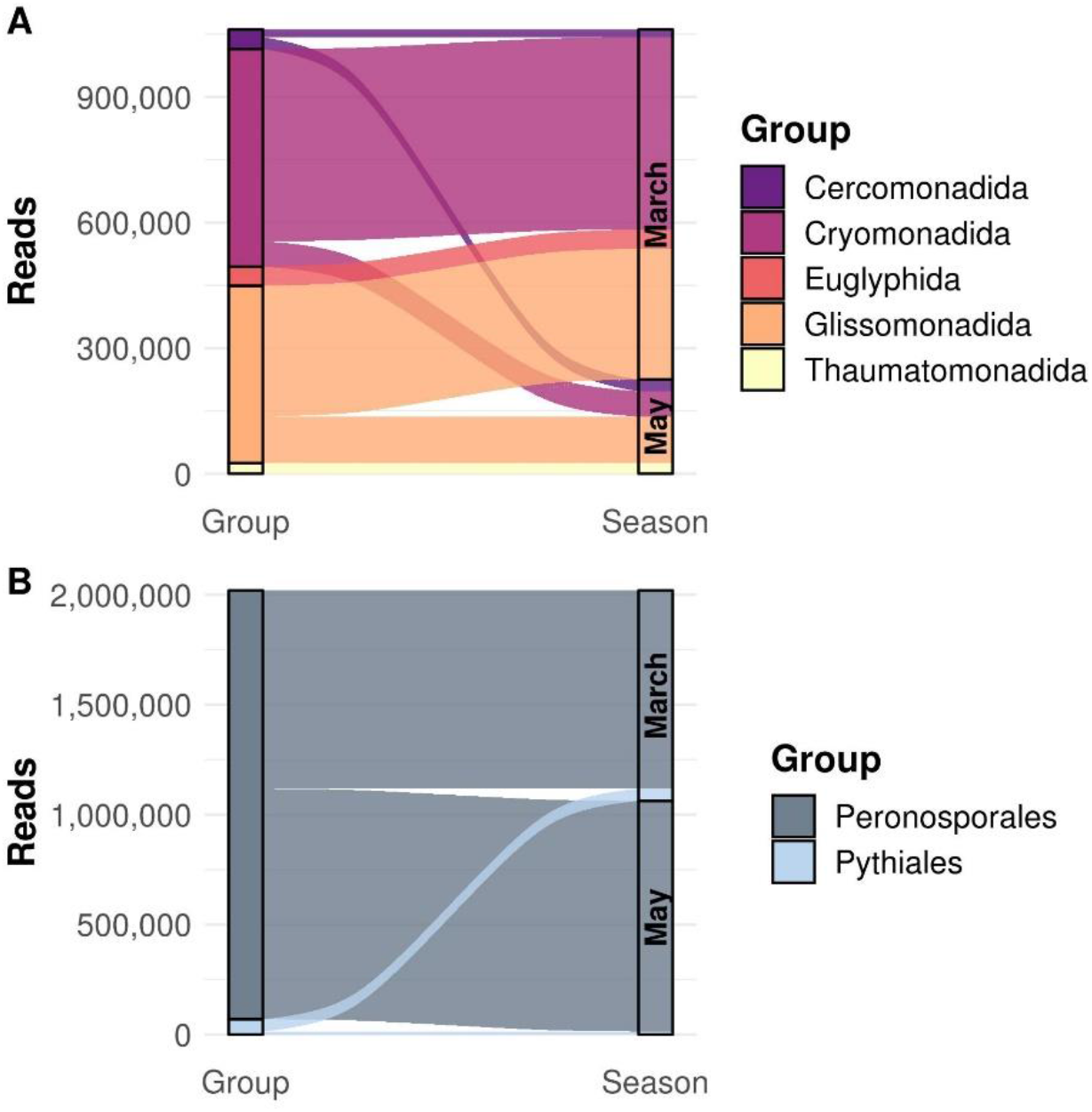
Sankey distribution diagram of cercozoan (A) and oomycete (B) orders in March and May samples. Orders represented by less than 1% of all reads were removed from the visualisation for the sake of clarity.

## 4. Discussion

In a recent study, Jauss et al. (2020) quantified the diversity of Oomycota and Cercozoa in canopy microhabitats and in litter and soil on the ground of the same floodplain forest. This allows for a direct comparison of the total diversity of these protists in the forest stand with the potential diversity of taxa distributed by air during two time points. We detected 122 and 81 OTUs of Cercozoa and Oomycota in air samples, respectively, which corresponds to 22 and 24% of the former reported total diversity of these protistan phyla. The high temporal variation, also reflected by the number of shared OTUs (Figure 4), suggests protistan distribution to be not restricted by dispersal-limitation, but rather indicates a continuous propagule rain of potentially invasive species and their accumulating resting stages occupying vacant niches. The vertical distribution of protists in air samples was rather homogeneous and did not differ between tree canopy and ground. In contrast, Jauss et al. (2020) found clear spatial patterns of oomycetes and cercozoans in tree canopies compared to the forest floor, suggesting that only part of the wind-borne propagule rain finds suitable conditions for survival in tree crowns due to habitat filtering. The temporal variation could be either related to temporal variations in the activity and distribution of protists, or more likely due to a dependency on the weather during the sampling. In March, the conditions were less favorable, with comparatively low temperatures and humidity with immediate previous precipitation events that have to be taken into account (Table 1). The remaining ground moisture might have prevented the lofting of protists through wind currents, while in May the atmospheric conditions were more preferable with higher average temperatures and a higher humidity. The conditions in May probably favored the lofting of protists into the atmosphere and their long-distance dispersal, leading to a higher protistan diversity and OTU richness (Figure 1), even though the wind speed was slower compared to March (Table 1). As wind speed was determined to be an important factor governing the species richness of microorganisms in air samples (Rogerson & Detwiler, 1999; Genitsaris et al., 2014), faster wind speeds in May probably could have revealed more protists. This suggests not a single factor, but rather the interplay between atmospheric conditions driving the species richness and community assembly in the air, while our samples possibly only represent the lower counts of what can be dispersed by air.

Wind dispersal is an important means for the distribution of microbial plant pathogens, and oomycetes are no exception (Fawke et al., 2015; Lang-Yona et al., 2018). Yet, comprehensive assessments of their abundances within the forest air were lacking. The presence of ~54% of all oomycete OTUs in both sampling events (Figure 4B) indicates a continuous presence of both peronosporean and pythialean oomycete spores and consequently a high proportion of potentially physiologically active oomycetes, including potential pathogens within forest ecosystems. Oomycetes pose a serious threat to forest health and functioning, it is therefore crucial to better understand their diversity and distribution patterns of the total forest ecosystem, including air samples (Derevnina et al., 2016; Ajchler et al., 2017; Jung et al., 2018; Lang-Yona et al., 2018). We detected no OTUs assigned to the orders Saprolegniales, Lagenidiales or Myzocytiopsidales, even though Jauss et al. (2020) found them in canopy and ground samples. All three orders are capable of forming dispersal stages, while their absence in our air samples could be due to a different timing of their sporulation, as our samples can only represent a snapshot of aerobic diversity.

Three dominant cercozoan orders were detected in air samples, but surprisingly no plant parasites of the Endomyxa. Testate amoebae from the orders Cryomonadida and Euglyphida occurred in high numbers. Cryomonadida (Thecofilosea) are filose amoeba with a robust extracellular organic tests (Adl et al., 2019). OTUs assigned to the Rhogostoma-lineage within the Cryomonadida dominated the samples. *Rhogostoma* species form resting stages resistant against desiccation for up to three months, although they form no cysts or zoospores (Mylnikova & Mylnikov, 2012; Öztoprak et al., 2020). *Assulina seminulum*, has a silica test with a remarkable size of 60-90 μm (Lara et al., 2010), and dominated the air dispersed Euglyphids, demonstrating that protists of this size can be still easily dispersed by air (Finlay, 2002). Not surprising was the dominance of Glissomonadida, represented by small flagellates of the families Sandonidae and Allapsidae. Their high abundance is consistent with observations Ploch et al (2016) and Jauss et al (2020). All these orders are an integral part of the protist phyllosphere microbiome (Agler et al., 2016; Dumack et al., 2017; Flues et al., 2018). Overall, their presence in the microbiome as well as their high abundance in air samples indicates canopies and their phyllosphere to be a potential filter not only for dust and particles (Weber et al., 2014; Chen et al., 2017), but also for microorganisms and potential plant pathogens.

## Conclusion

A significant temporal variation in oomycetes indicates protistan community and, correspondingly, pathogen assembly to be driven by random factors and neutral processes, while spatial differences in the vertical distribution of cercozoans and oomycetes were not found. Accordingly, wind dispersal alone may well explain the ubiquitous presence of Cercozoa and Oomycota (and likely of other protistan taxa) in the floodplain forest. Our results further contribute to the understanding of how protists disperse, and which factors drive the distribution of plant pathogens within forest ecosystems.

## Acknowledgements

This paper was supported by the Priority Program SPP 1991: Taxon-omics − New Approaches for Discovering and Naming Biodiversity of the German Research Foundation (DFG) with funding to MB (1907/19-1) and MS (Schl 229/20-1). The authors would like to thank Rolf Engelmann for his assistance with the field work by operating the canopy crane, and the German Centre for Integrative Biodiversity Research (iDiv) for providing the site access.

## Author Contributions

MS and MB conceived the study, RW and StS designed the sampling. BS assisted the library preparation. SW assisted the sampling and contributed valuably to the discussion. AN performed the sampling, laboratory work, bioinformatics analyses and outline of the manuscript. R-TJ supervised the study, sampling and bioinformatic analyses, and wrote the manuscript. All authors contributed to the manuscript and approved the final version.

## Data availability

Raw sequence data have been submitted to the European Nucleotide Archive (ENA) database under the Bioproject number PRJEB37525, with accession numbers ERS5388855 (Cercozoa) and ERS5388854 (Oomycota) respectively.

All figures, codes and detailed bioinformatic/statistical methods used in this study are available at https://github.com/RJauss/ToTheCanopyAndBeyond.

